# Urinary biochemical ecology reveals microbiome-metabolite interactions and metabolic markers of recurrent urinary tract infection

**DOI:** 10.1101/2024.10.22.619727

**Authors:** Michael L. Neugent, Neha V. Hulyalkar, Debasish Ghosh, Ceejay N. Saenz, Philippe E. Zimmern, Vladimir Shulaev, Nicole J. De Nisco

**Affiliations:** Department of Biological Sciences, The University of Texas at Dallas, Richardson, Texas; Department of Urology, The University of Texas Southwestern Medical Center, Dallas, Texas; Department of Biological Sciences, The University of North Texas, Denton, Texas

**Keywords:** urinary microbiome, metabolomics, ecology, urinary tract infection, biomarker

## Abstract

Recurrent urinary tract infections (rUTIs) are a major clinical challenge and their increasing prevalence underscores the need to define host-microbiome interactions underlying susceptibility. How the urinary microbiota engages with the biochemical environment of the urogenital tract is yet to be fully defined. Here, we leverage paired metagenomic and quantitative metabolomic data to establish a microbe-metabolite association network of the female urinary microbiome and define metabolic signatures of rUTI. We observe unique metabolic networks of uropathogens and uroprotective species, highlighting potential metabolite-driven ecological shifts influencing rUTI susceptibility. We find distinct metabolites are associated with urinary microbiome diversity and identify a lipid signature of active rUTI that accurately distinguishes cases from controls. Finally, we identify deoxycholic acid as a prognostic indicator for UTI recurrence. Together these findings provide insight into microbiome-metabolite interactions within the female urinary tract and highlight new biomarkers for the development of new diagnostic tools to improve patient outcomes.

## Introduction

Urinary tract infections (UTIs) are among the most common bacterial infections in adults worldwide with many UTIs progressing into recurrent UTI (rUTI) or chronic UTI.^1,2^ Frequent UTI recurrence, despite antibiotic intervention, suggests underlying alterations in the urogenital ecosystem, including both host and microbial factors, contribute to susceptibility.^3–5^ Indeed, the decline in circulating estrogen levels associated with menopause is one of the primary risk factors for rUTI with reports of up to 50% of UTIs progressing to rUTI in postmenopausal women.^1,2,5,6^ Treatments for rUTI have focused on modifying the microbial environment of the urinary tract, often through the goal of achieving sterility via long-term and prophylactic antibiotic regimens.^4,7,8^ However, the discovery of a urinary microbiome in healthy individuals has highlighted the need to develop new treatment paradigms for rUTI focused on eradicating pathogens while restoring a healthy urinary microbiome.^9,10^

The microbial communities of the urogenital tract, here termed the urinary microbiome, have been extensively characterized and linked to both health and disease.^10–22^ Research has shown that, in females, urinary microbial communities dominated by *Lactobacillus* species are linked to health, while those with *Escherichia coli* are often associated with UTI.^5,23^ Despite extensive compositional profiling of the urinary microbiome, less is known about how these microbial communities interact with the metabolic environment of the urine. The biochemical landscape of the urinary tract may reflect key physiological and pathological states.^24–26^ The urinary metabolome, in particular, could reveal host responses to infection and metabolic relationships between the host and the urinary microbiota.^27^ Investigating the taxonomic and biochemical ecology of the urinary microbiome may also facilitate the identification of biomarkers to improve UTI diagnosis, prognosis, and treatment. Previous studies have suggested links between urinary metabolites like putrescine and uropathogenic *E. coli* (UPEC), but the urinary metabolome and its relationship with the taxonomic ecology of the urinary microbiome remains largely unexplored.^28^

The concept of “biochemical ecology” refers to the functional interactions between metabolites and microbial communities. Emerging evidence suggests that metabolites not only result from microbial activity but also shape the microbial ecosystem itself, potentially influencing microbiome function and susceptibility to infection.^29–31^ However, little is known about the relationship between the taxonomic and biochemical ecology of the urinary tract and how these may jointly contribute to the pathophysiology of rUTI. Building on our previous reports from a cross-sectional cohort of postmenopausal women designed to study UTI, this study integrates metagenomic and metabolomic data to establish associations between urinary biochemical and taxonomic ecology, and to identify metabolic signatures associated with rUTI. By linking urinary metabolic profiles to microbial ecology, we aim to provide a more comprehensive understanding of the biochemical factors influencing urinary tract health and infection susceptibility.^5^

Specifically, we investigate whether distinct biochemical signatures are linked to microbial community structure and disease states using a multiplexed targeted metabolomics assay quantifying more than 600 metabolites.^32,33^ Our findings suggest that the urinary metabolome reflects not just microbial activity but also broader host responses, such as oxidative stress, potentially linking local inflammation to microbial compositional diversity. In addition to defining new and distinct associations between urinary metabolites and different microbiome species, we identified a suite of urinary lipids that exhibit potential as accurate diagnostic biomarkers of active UTI. We also found that urinary deoxycholic acid was prognostic for increased rUTI risk and associated with a significant decrease in recurrence-free days. Together, these insights reveal critical links between biochemical and microbial ecologies in the female urinary tract, offering new avenues for precision diagnostics that may lay the foundation for improved patient outcomes in the clinical management of rUTI.

## Results

### Targeted LC-MS/MS of the urinary polar metabolome and lipidome

To quantitatively characterize the urinary metabolome associated with recurrent urinary tract infection (rUTI) and the urinary microbiome, we performed targeted liquid chromatography tandem mass spectrometry (LC-MS/MS)-based metabolite profiling using the Biocrates MxP® Quant 500 assay^34^ of more than 600 polar and non-polar metabolites in urine collected from a previously published cross-sectional cohort of postmenopausal women we designed to study recurrent UTI.^5^ This cohort consists of three groups of postmenopausal women differentiated by their UTI history and current UTI status (*n*=75, 25 per group) (**Fig. 1A**).^5^ Group 1 (No UTI History) has no lifetime history of UTI and was used as a healthy postmenopausal comparator. Groups 2 and 3 consisted of postmenopausal women with a previous history of rUTI but differed on UTI status at time of urine donation. Group 2 (rUTI History, UTI(-)) consisted of women with a recent history of rUTI who were not experiencing a UTI at the time of urine donation, while Group 3 (rUTI History, UTI(+)) consisted of postmenopausal women with a clinically confirmed UTI at the time of urine donation.^5^

**Figure 1.**
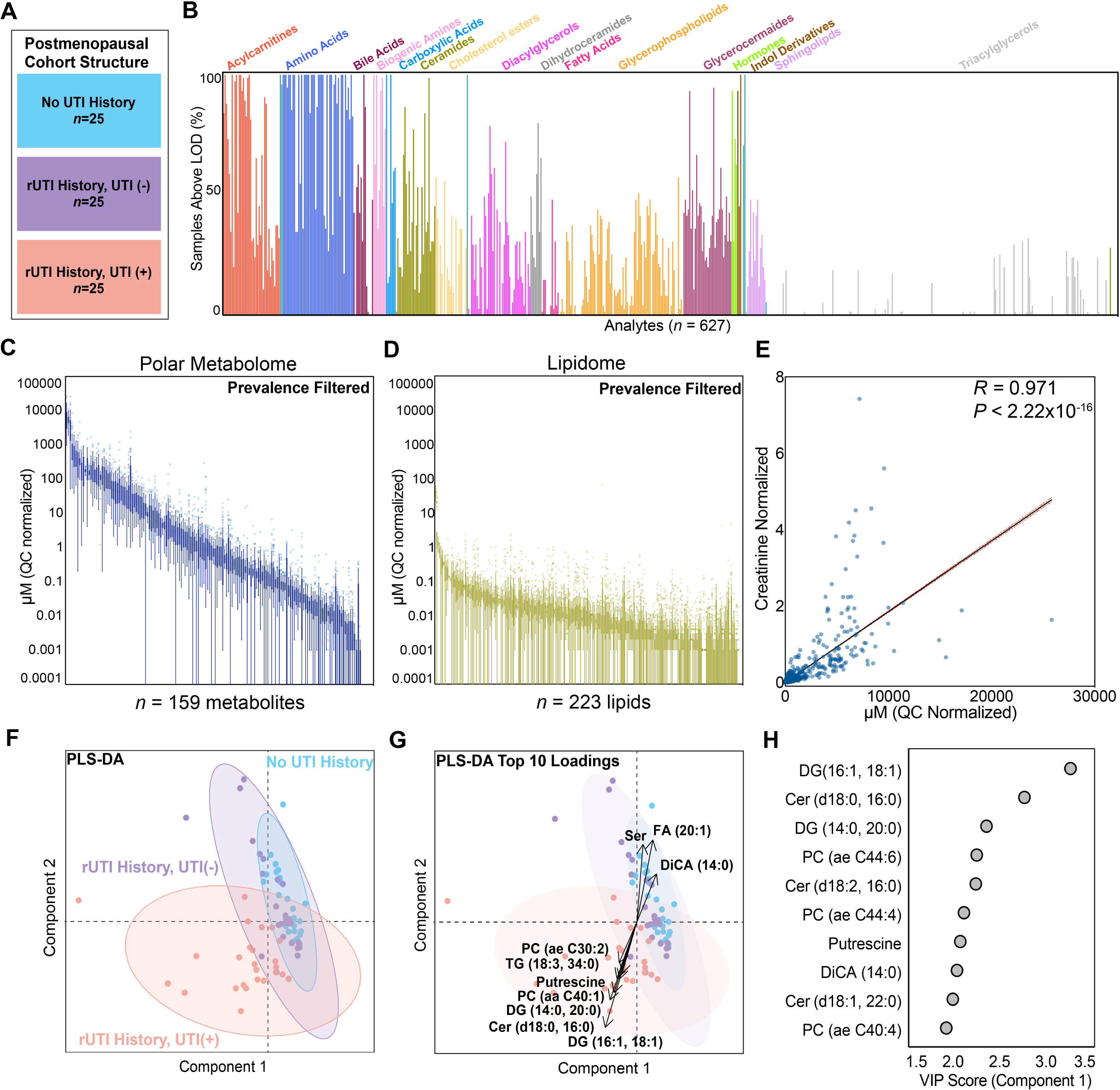
Targeted LC-MS/MS of the urinary polar metabolome and lipidome. (A) Cohort structure breakdown. (B) Overview of metabolite classes detected by metabolomic assay. Y-axis refers to the proportion of the cohort where a given metabolite was detected above the limit of detection (LOD). (C) Polar metabolites retained after filtering. (D) Non-polar metabolites retained after filtering. (E) Correlation of raw and creatinine normalized metabolite abundances. *P-*value generated by permutation. (F) PLS-DA ordination plot of the cohort samples. 95% confidence ellipses are provided to estimate group ordination. (G) PLS-DA biplot overlaying the top 10 loading contributing to the PLS-DA group ordination. (H) Top 10 VIP scoring metabolites.

The urinary metabolites measured included a broad range of compounds from polar to non-polar metabolites (**Fig. 1B**). A total of 159 polar metabolites and 223 lipid species were selected for further downstream analysis (**Fig. 1C, D; Table S1**). We next assessed correlation between metabolite concentrations and the creatinine-normalized abundances (*R* = 0.971, *P* < 2.22×10^-16^, *n* = 28,350 pairwise comparisons) (**Fig. 1E**). While both values have utility for distinct questions, we chose to focus our analysis on the raw urinary concentration of each metabolite given our intention to associate the biochemical ecology with the local microbial ecology, a set of variables not necessarily associated with glomerular filtration and creatinine excretion rate.^3,35,36^

To evaluate large-scale differences between groups and identify metabolites most associated with group identity, we performed Partial Least Squares Discriminant Analysis (PLS-DA). This analysis revealed distinct clustering of samples from the rUTI History, UTI(+) group, while samples from the rUTI History, UTI(-) and No UTI History groups clustered together (**Fig. 1F, G**). Additionally, the top 10 metabolites driving group separation, as indicated by the highest Variable Importance Projection (VIP) scores, were predominantly lipid species, including the diacylglycerol DG(16:1/18:1) and the ceramide Cer(d18:0/16:0) (**Fig. 1H, Fig. S1**). Interestingly, putrescine, a biogenic amine produced by UPEC during infection, was among the most important metabolites for establishing group identity.^28,37^

### Urine biochemical ecology is moderately associated with the taxonomic ecology

To characterize the association between metabolic and microbial ecological signatures defining rUTI susceptibility and active UTI, we tested the hypothesis that there may be broad associations between urinary microbiome and biochemical ecology. We observed no difference in the biochemical α-diversity between groups even though there was a detectable difference in microbial α-diversity between the rUTI History, UTI(-) and rUTI History, UTI(+) groups (**Fig. 2A, B; Table S2**).^5,38^ The taxonomic α-diversity also exhibited a much larger range (1.003 - 22.05) than that of the biochemical ecology (2.26-7.15) (**Fig. 2B**). Additionally, we observed no association between the overall biochemical and taxonomic α-diversity of urine (**Fig. 2C**). These data suggest that while the urinary microbiome and metabolome are components of the same ecosystem, their α-diversities do not appear to be associated in the cohort studied. However, we observed significant associations between individual metabolites and taxonomic α-diversity. Urinary concentrations of methionine sulfoxide (Met-SO), citrulline (Cit), and trimethylamine oxide (TMAO) were all significantly positively associated with both the Shannon and Simpson taxonomic α-diversity (**Fig. 2D-F**).

**Figure 2.**
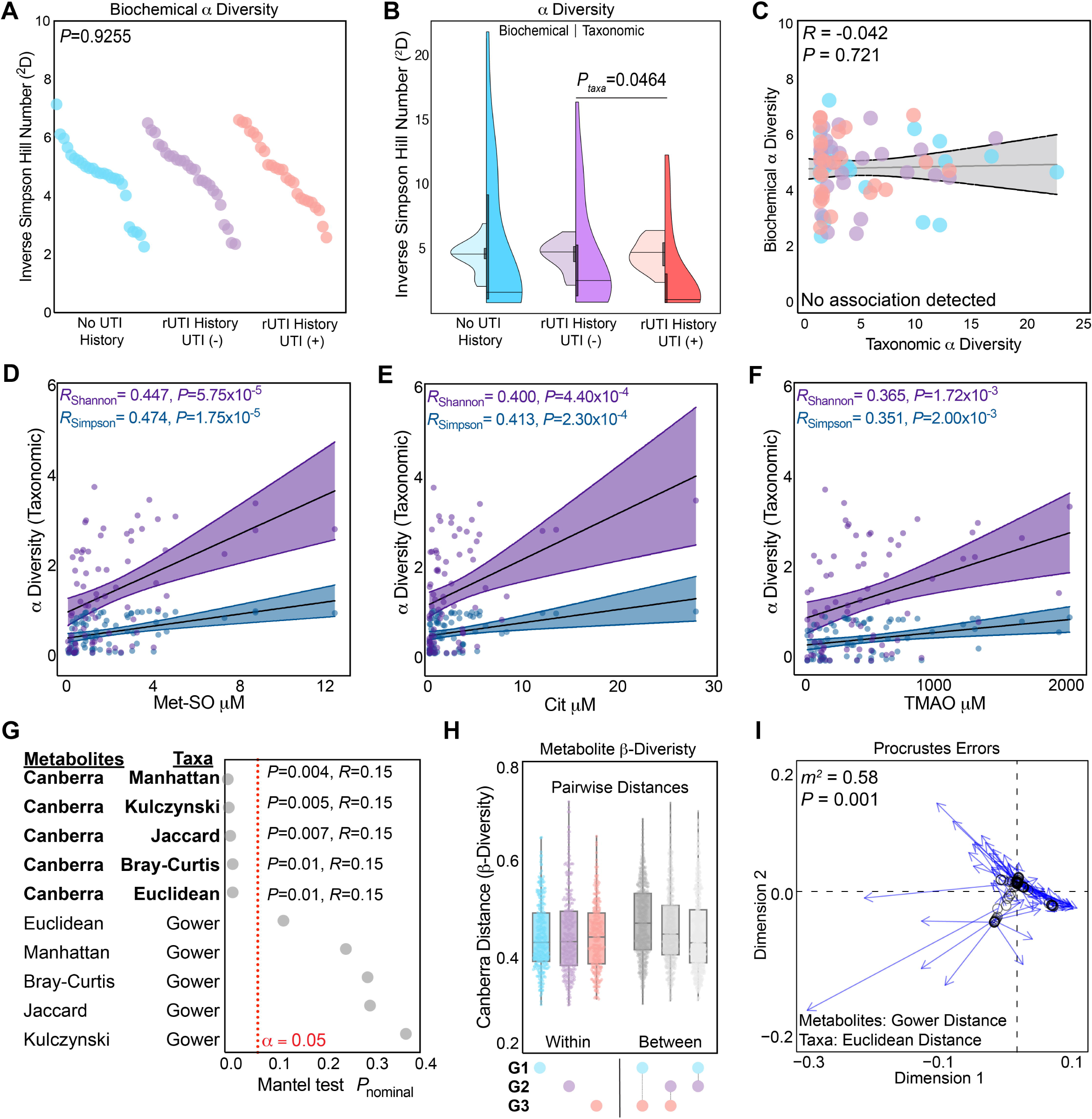
Urinary biochemical ecology is moderately associated with the taxonomic ecology. (A) Distribution of α diversity (Inverse Simpson Hill Number) among the three cohort groups (*n*=75 total, *n*=25 per group). *P*-value generate by Kruskal-Wallis nonparametric ANOVA. (B) Comparison of the distribution of biochemical (left) and taxonomic (right) α diversity (Inverse Simpson Hill Number). Violin is drawn from minimum to maximum of the distribution. Median is denoted by a line and boxes are drawn to represent the interquartile range. (C) Spearman correlation of biochemical and taxonomic α diversity. *P*-value generated by permutation. 95% confidence bands border the linear regression line. (D-F) Spearman correlation of taxonomic α diversity with the urinary concentration of individual metabolites, methionine sulfoxide (Met-SO) (D), citrulline (E), and trimethylamine oxide (F) by both taxonomic Shannon and Simpson metrics. *P*-value generated by permutation. 95% confidence bands border the linear regression line. (G) Mantel Test screening of the association between biochemical and taxonomic β diversity combinations. (H) Analysis of pairwise Canberra β diversity distances within and between cohort groups. Boxes draw to represent the median and interquartile range. Whiskers drawn to the minima and maxima of the data distribution. (I) Procrustes errors from analysis of the association between biochemical and taxonomic β diversity. For Procrustes analysis, PCoA ordination of the biochemical β diversity was used as the target, while taxonomic β diversity ordination was rotated and scaled. *P*-value generated by permutation Protest.

We next hypothesized that urine samples with distinct taxonomic compositions would also have distinct metabolite compositions. To test this hypothesis, we analyzed covariance of β-diversity between the taxonomic compositional data and the metabolomic data (**Fig. 2G, H, I**). For both datasets, we screened a number of commonly used methods to calculate β-diversity distance, including the Bray-Curtis, Jaccard, Euclidean, Manhattan, Canberra, Gower, and Kulczynski.^39–41^ To measure covariance between all combinations of the distances of taxonomic and biochemical ecology, we used the Spearman variant of the Mantel test.^42^ We observed significant covariance (*P* < 0.05) when the biochemical β-diversity was calculated using the Canberra distance (**Fig. 2G**). However, the association detected was relatively weak (*R* = 0.15). Canberra distance normalizes feature differences, highlighting the potential role of low-abundance metabolites in linking taxonomic and biochemical β-diversity.

We further observed that pairwise biochemical β-diversity distances were higher when comparing between disease groups relative to within, suggesting that differences in disease state are concurrently associated with altered urinary metabolome (**Fig. 2H**). Between group metabolic β-diversity differences were largest between Group 1 (No UTI History) and Group 3 (rUTI History, UTI (+)) (*P_Wilcoxon_*< 10^-9^). Given the limitations of pairwise tests like the Mantel test in detecting more complex associations, we used Procrustes analysis to assess the alignment of independent biochemical and taxonomic PCoA ordinations.^43^ This analysis found most significant, moderate alignment (*m^2^* = 0.58, *P* = 0.001) when using the Gower distance for metabolites and the Euclidean distance for taxa (**Fig. 2I**). Similarly to as observed in Fig. 2G, associations between urinary biochemical and taxonomic β-diversity were identified when modeling biochemical distances using metrics that normalize each feature to its total range across all samples. Taken together, these data suggest the covariance between urinary taxonomic and biochemical β-diversity is driven primarily by low abundance metabolites.

### *Escherichia* is a metabolic hub defined by a high degree of unique metabolite associations

We next sought to define the structure of the individual taxon-metabolite association network. To accomplish this, we performed an all-against-all Spearman correlation analysis between the urinary microbiome genus-level taxonomic profile and urinary metabolomic profile. For this analysis we focused solely on Groups 1 and 3 and removed the intermediate disease state Group 2 so that clear comparisons could be made between cases and controls. We used the significant (*R* > |0.3|, *P* < 0.05) correlation results between taxa-metabolite pairs to generate both positive and negative association networks with nodes representing metabolites (purple) or taxa (pink) and edges drawn only between taxa and metabolites (**Fig. 3A, B; Table S3**). The positive association network was relatively sparse with a striking separation between the larger metabolite-taxa interaction network and that of a single taxon, *Escherichia* (**Fig. 3A**). Not only was *Escherichia* associated with a highly distinct set of metabolites, but it was also the taxon node with highest degree, or number of first neighbors (*n* = 122 metabolites) (**Fig. 3A, C; Table S4**). Most metabolites associated with *Escherichia* abundance were not strongly connected to other taxa as indicated by a relatively low topological coefficient (*T(i)* = 0.0139) (**Fig. 3C**). These data suggest that urinary *Escherichia,* which in this dataset is largely associated with active UTI, is associated with a unique set of urinary metabolites, few of which are associated further with additional taxa.^5^ Among these metabolites were putrescine (*R* = 0.661, *P* = 1.37×10^-6^), one of the major polyamines produced by *E. coli,* and the diacylglycerol, DG(14:0, 20:0) (*R* = 0.638, *P* = 4.18×10^-6^).^28,37^ We hypothesize that this observation is evidence of a unique urinary metabolic signature associated with UTI, the majority of which in this cohort were culture-confirmed UPEC infections as we have previously reported.^5^

**Figure 3.**
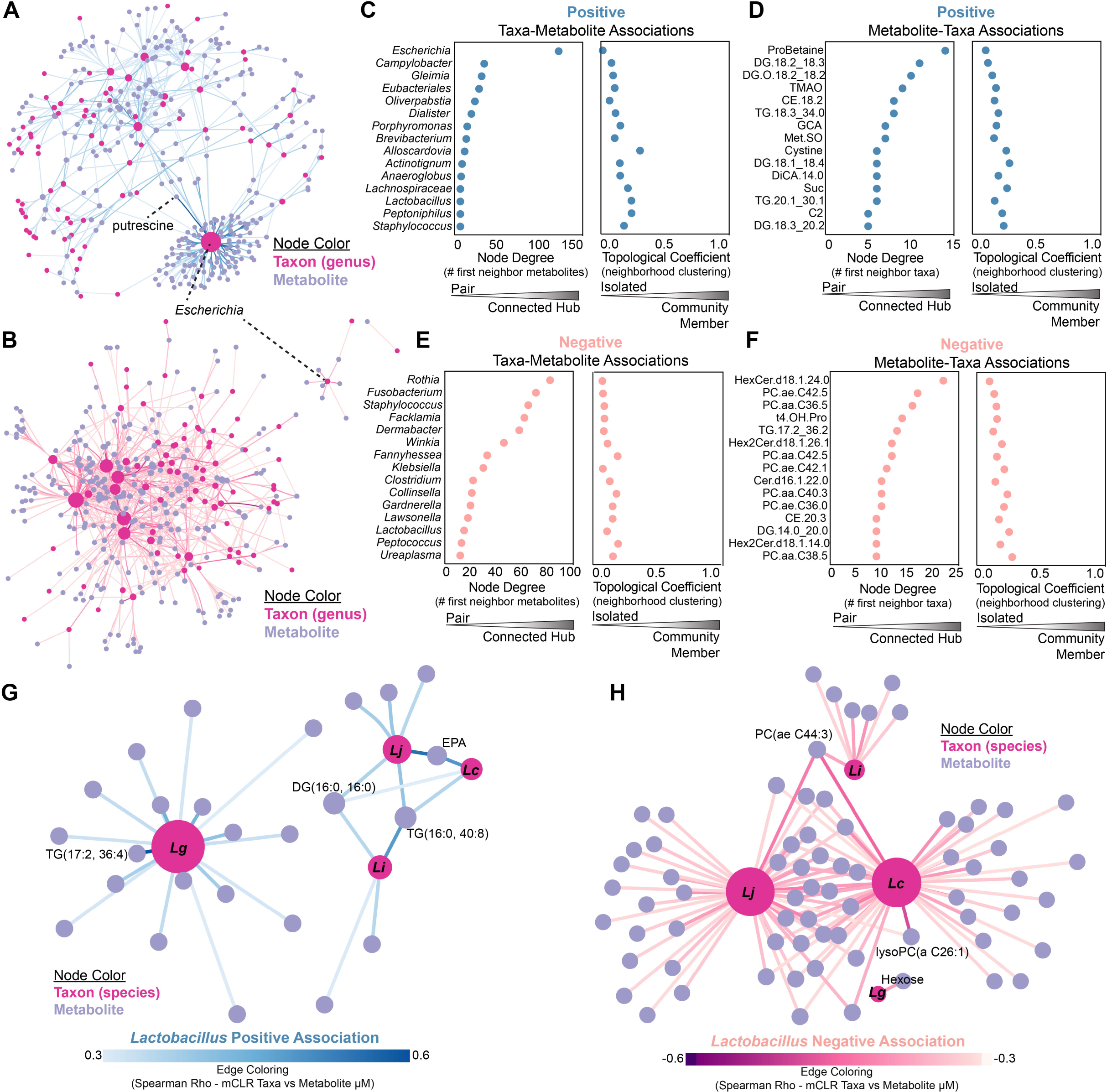
Taxa metabolite network analysis. (A) Network representation of significant (*R* > 0.3, *P* < 0.05) positive taxon (red nodes) metabolite (blue nodes) associations. (B) Network representation of significant (*R* < −0.3, *P* < 0.05) negative taxon (red nodes) metabolite (blue nodes) associations. Edges represent the Spearman Correlation between metabolite concentrations and mCLR transformed taxon relative abundance. Taxon node size is proportional to the number of first neighbor metabolites. (C) Network metrics, nodal degree and topological coefficient, of taxa in the positive association network. (D) Metabolite nodal degree and topological coefficient in the positive association network. (E) Taxon nodal degree and topological coefficient in the negative association network. (F) Nodal degree and topological coefficient of metabolites in the negative association network. Network representation of significant (G) positive (*R* > 0.3, *P* < 0.05) and negative (H) species level *Lactobacillus* (red nodes) metabolite (blue nodes) positive associations. Edges represent the Spearman Correlation between metabolite concentrations and mCLR transformed taxon relative abundance. Taxon node size is proportional to the number of first neighbor metabolites.

When analyzed from the perspective of the metabolites, proline betaine (ProBetaine), a proline derivative also known as stachydrine, was identified as the highest degree metabolite node, being connected to more taxa than any other metabolite (*n* = 14 genera) (**Fig. 3D**).^44^ ProBetaine was positively associated with the genera *Slackia* (*R* = 0.436), *Schaalia* (*R* = 0.398), *Peptococcus* (*R* = 0.383), and *Mobiluncus* (*R* = 0.367). The negative taxa-metabolite association network was more densely connected with the genera, *Rothia* (*n* = 82 metabolites), *Fusobacterium* (*n* = 71 metabolites), *Staphylococcus* (*n* = 65 metabolites), *Facklamia* (*n* = 62 metabolites), and *Dermabacter* (*n* = 58 metabolites) making up the top 5 taxa connected with the greatest number of metabolites (**Fig. 3B, E; Table S5**). The hexosylceramide, HexCer(d18:1/24:0), had the highest degree, being negatively associated with 22 genera (**Fig. 3F**).

### The unique metabolite network of *Lactobacillus gasseri* separates it from other urogenital lactobacilli

While much of the taxa-metabolite network analysis revealed associations between metabolites and known uropathogens, we further sought to determine the *Lactobacillus*- metabolite associations to identify biochemical features associated with these beneficial microbial communities.^45^ Our analysis of the species-level *Lactobacillus* positive association network revealed two non-interacting networks – one composed solely of *L. gasseri* and the other composed of *L. jensenii*, *L. crispatus*, and *L. iners* (**Fig. 3G**). *L. gasseri* positively associated with a unique and distinct set of 17 metabolites. The strongest metabolite association with *L. gasseri* was the triglyceride TG(17:2, 36:4), a triacyl glycerol (*R* = 0.542, *P* = 0.00018). The remaining *Lactobacillus* species all associated with common metabolites within a shared network neighborhood (**Fig. 3G**). Both *L. jensenii* and *L. crispatus* exhibited relatively strong associations with the fatty acid, eicosapentaenoic acid (EPA) (*R_jensenii_* = 0.503, *R_crispatus_* = 0.450), while all three taxa, *L. iners, L. jensenii,* and *L. crispatus* were correlated with the triacylglycerol, TG(16:0, 40:8) (*R_iners_* = 0.447, *R_jensenii_* = 0.377, *R_crispatus_* = 0.354). The diacylglycerol, DG(16:0, 16:0), was found to be associated with *L. jensenii* (*R* = 0.350) and *L.* iners (*R* = 0.342) and more weakly with *L. crispatus* (*R* = 0.301) (**Fig. 3G**). The negative association network also separated *L. gasseri* from the other urogenital *Lactobacillus* species and revealed that *L. jensenii* and *L. crispatus* were highly interconnected, negatively correlating with a common set of 28 metabolites. The strongest shared anti-correlation of *L. crispatus* and *L. jensenii* was the phospholipid, PC(ae C44:3) (*R_crispatus_* = −0.426, *R_jensenii_* = −0.402), which was also negatively associated with *L. iners* (*R* = −0.359) (**Fig. 3H**). *L. gasseri* was negatively associated with only a single metabolite, urinary hexose (**Fig. 3H**). Given the highly similar chromatographic behavior and mass spectral features of hexoses, such as glucose, galactose, and fructose, we cannot robustly determine which hexose(s) drive this negative metabolite association.^46^ Together, our data suggests that urinary *L. gasseri* communities are both positively and negatively associated with a urinary metabolic signature that is unique among the major urogenital lactobacilli.

### Active UTI is associated with a distinct metabolic signature

We hypothesized that the urinary metabolome between PM women who do not get UTIs and those actively presenting with a UTI would be significantly different and may harbor novel biomarkers of active UTI. To test this hypothesis, we performed a differential enrichment analysis screen of metabolites between Group 1 (No UTI History) and Group 3 (rUTI History, UTI(+)) using the Wilcoxon rank sum test (**Fig. 4A, Table S6**). This analysis identified many significantly differentially abundant metabolites enriched either in the No UTI History (*n* = 9 metabolites) or rUTI History, UTI(+) (*n* = 149 metabolites) (**Fig. 4B, C**). The observation that putrescine was among the most differentially abundant metabolites enriched in urine during active rUTI (*FC_median_*= 12.63, *Q* = 3.77×10^-6^) is in line with expectations based on previous observations and can be attributed largely to *Escherichia coli* as putrescine was found to be highly correlated to *Escherichia* in our taxa-metabolite network analysis in Figure 3 (**Fig. 4B, D**).^28^ The remaining top 10 differentially abundant metabolites enriched during active rUTI were all lipid species, such as diacylglycerols (DG), ceramides (Cer), and glyceroceramides (Hex2Cer) (**Fig. 4B, E, F, G**).

**Figure 4.**
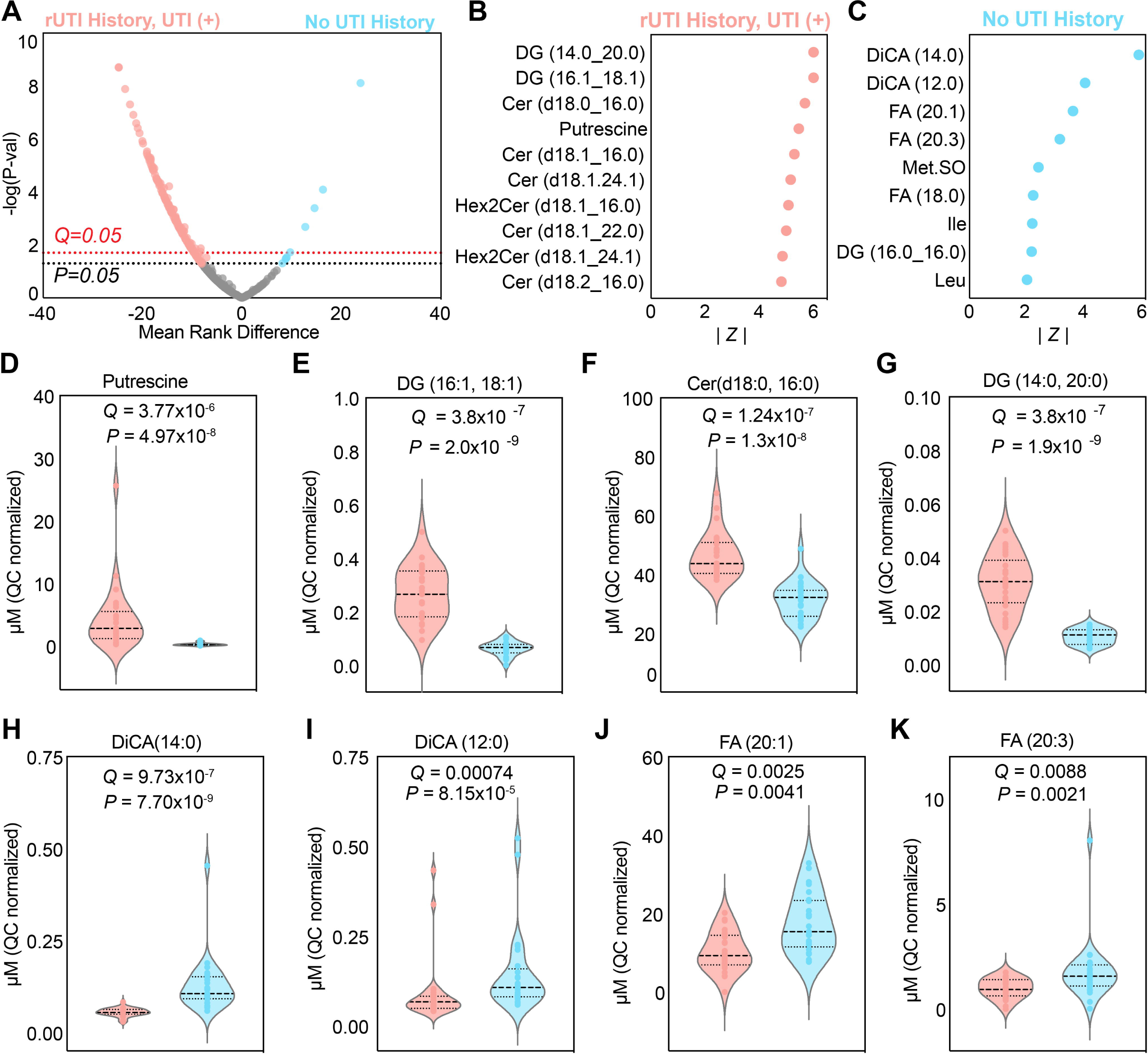
Active UTI is associated with distinct urine metabolic signature. (A) Differential enrichment analysis of urinary metabolites between the No UTI History (Blue) and rUTI History, UTI(+) (Red) groups. *P*-value generated by the Wilcoxon Rank Sum test. *Q*-value generated by false discovery rate. Absolute value of the Wilcoxon Rank statistic of the top 10 metabolites differentially enriched in the (B) rUTI History, UTI (+) group and in the (C) No UTI History group. (D-G) Comparison of the distributions for individual metabolites differentially enriched in the rUTI History, UTI (+) group. (H-K) Comparison of the distributions for individual metabolites differentially enriched in No UTI History group. *P*-value generated by the Wilcoxon Rank Sum test. *Q*-value generated by false discovery rate.

In contrast, we observed a separate set of 9 metabolites significantly enriched in the urine of PM women in Group 1, the women with No UTI history. Differentially abundant metabolites include three fatty acids (FA 20:1, 20:3, and 18:0), methionine sulfoxide (MetSO), the amino acids isoleucine (Ile) and leucine (Leu), and a diacylglycerol (DG) (**Fig. 4C, H, I, J, K**). Interestingly, two dicarboxylic acid metabolites, DiCA(14:0) and DiCA(12:0), were the most significant differentially abundant metabolites enrich in PM women who do not get UTI (*Q* < 10^-7^, *Q*=0.00074) (**Fig 4 C, H, I)**. These metabolites have previously been detected in urine and are typically linked to gamma-oxidation of long-chain fatty acids.^47^ To further explore these findings, we compared metabolite profiles between the No UTI History group and the rUTI History, UTI (-) group. This analysis confirmed that DiCA (14:0) and DiCA (12:0) were also significantly enriched in the No UTI History group (**Fig. S2**), reinforcing their potential role in differentiating metabolic signatures associated with UTI susceptibility. While, in many cases, we cannot robustly decouple host-derived metabolites from microbial metabolites, these data do suggest a unique urinary metabolic signature associated with women who do not experience UTI.

### Taxa associated with metabolites differentially enriched in No UTI history

Since the distinct set of metabolites enriched in women with no UTI history may be linked to urogenital health, we next investigated the taxonomic associations between these metabolites and the urinary microbiome. We performed a Spearman correlation analysis between the quantitative abundance of the nine metabolites enriched in the No UTI History group and the urinary microbiome taxonomic profile identified by MetaPhlAn4 in the same group.^48^ This analysis revealed 84 taxa-metabolite associations passing at least nominal significance at the genus and species levels (**Figure 5A, B, Table S7**). Of note, we observed robust associations between methionine sulfoxide (MetSO) with 4 genera and 12 species. The genus, *Lactobacillus*, as well as the species *Lactobacillus crispatus*, were the most negatively correlated taxa with urinary MetSO concentration (*R_Lactobacillus_* = −0.526, *R_L.crispatus_*= −0.551) (**Fig. 5C, D**). Conversely, we observed a relatively strong positive association between MetSO and the dysbiotic taxon, *Streptococcus anginosus* (*R* = 0.550, *P* = 0.004) (**Fig. 5E**). The genus, *Ureaplasma*, which is considered an opportunistic pathogen in the female urogenital tract, was positively associated with DiCA (12:0) (*R* = 0.469, *P* = 0.018) (**Fig. 5A**).^49^ We further observed a modest positive association between urinary fatty acid (18:0) and *Staphylococcus epidermidis* (*R* = 0.460, *P* = 0.012) (**Fig. 5F**).

**Figure 5.**
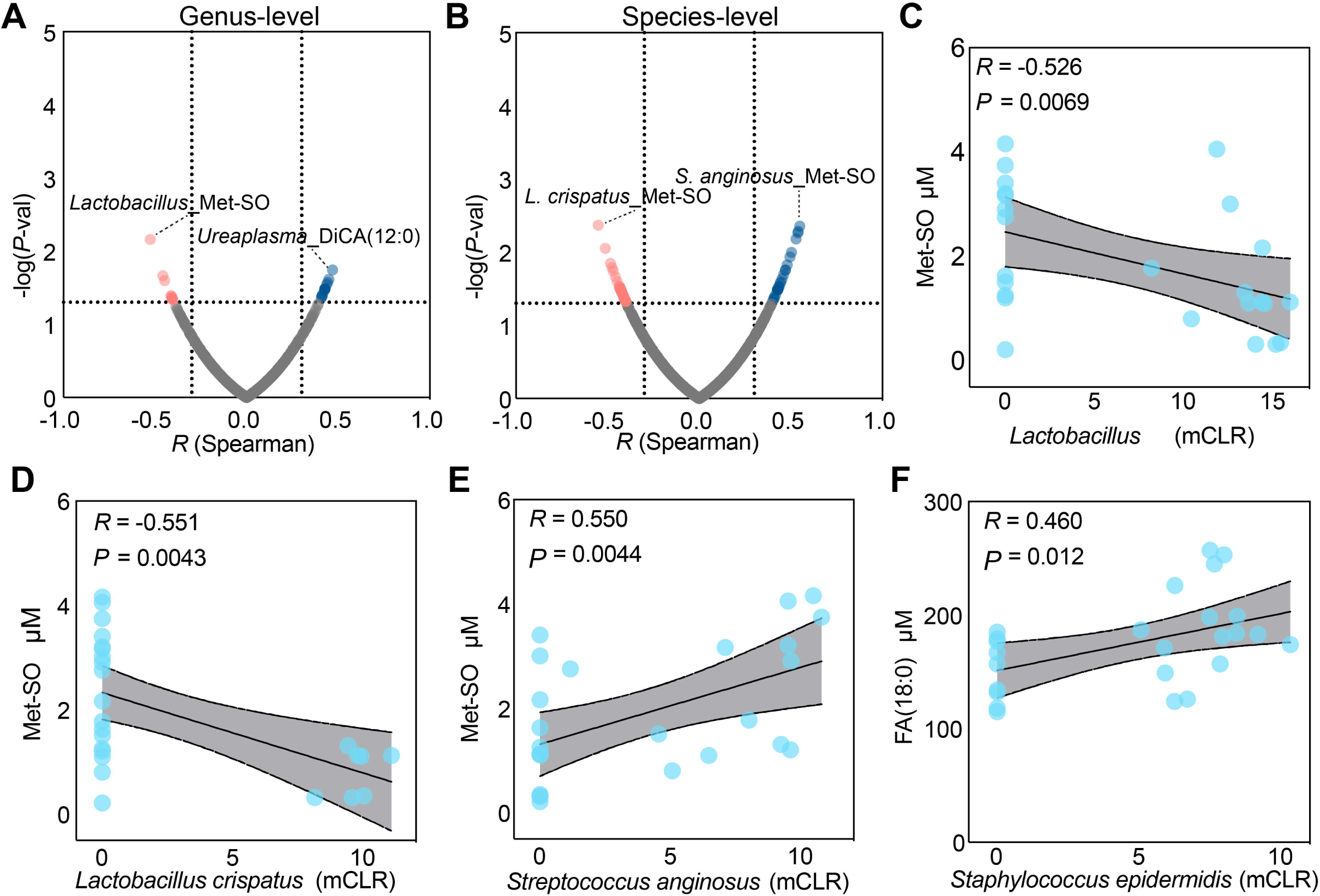
Taxa associated with urinary metabolites differentially enriched in women with No UTI history. Spearman correlation analysis between urinary metabolites differentially enriched in the No UTI History cohort and the urinary microbiome taxonomic profile at the (A) genus and (B) species level. (C-F) Scatter plots of significant taxa-metabolite correlations. *P*-value generated by permutation.

### Urinary lipids as biomarkers for active rUTI

Given our observation that many of the differentially abundant metabolites enriched in women with active rUTI were lipids (**Fig. 4B**) and that lipids contributed the most to group identity by PLS-DA (**Fig. 1E-G**), we hypothesized the urinary lipids may be an indicator or perhaps novel biomarker of active rUTI. When summed, the total assayed lipid content detected in urine exhibited a modest elevation in active rUTI (Group 3) as compared to the combined UTI(-) samples from Groups 1 and 2 (*FC_median_* = 1.32, *P* = 0.00023) (**Fig. 6A**). We then hypothesized that a specific subset of urinary lipids could differentiate case (active rUTI) from control (combined rUTI history, UTI(-) and No UTI history) groups. To test this hypothesis, we performed elastic net regularization on assayed lipids to identify an optimal subset of lipids distinguishing active rUTI cases from controls. This analysis identified 11 lipids in an optimal multivariate model whose abundance was able to distinguish the active rUTI samples in the rUTI History, UTI(+) group from the control samples in the combined rUTI History, UTI(-) and No UTI History group (**Fig. 6B, Fig. S3**). Diacylglycerol (16:1, 18:1), which was also the second most differentially abundant metabolite enriched in the UTI(+) group contributed the most to the multivariate model with a feature importance of 0.98 (**Fig. 6B**). To further assess the predictive power of this set of discriminating lipids, we performed logistic regression and leave one out cross validation (LOOCV) analysis (**Fig. 6C, D**). Logistic regression indicated that the 11 lipid metabolites were able to strongly discriminate cases from controls (*G^2^* = 72, *P* = 1.3×10^-11^) and the area under the receiver operator characteristic (ROC) curve of the 11 lipid metabolite multivariate model calculated by LOOCV was 0.9064 (**Fig. 6D**). These results suggest that this discriminating lipid signature has excellent sensitivity and specificity on cross validation; however, these results should be validated in an independent cohort.

**Figure 6.**
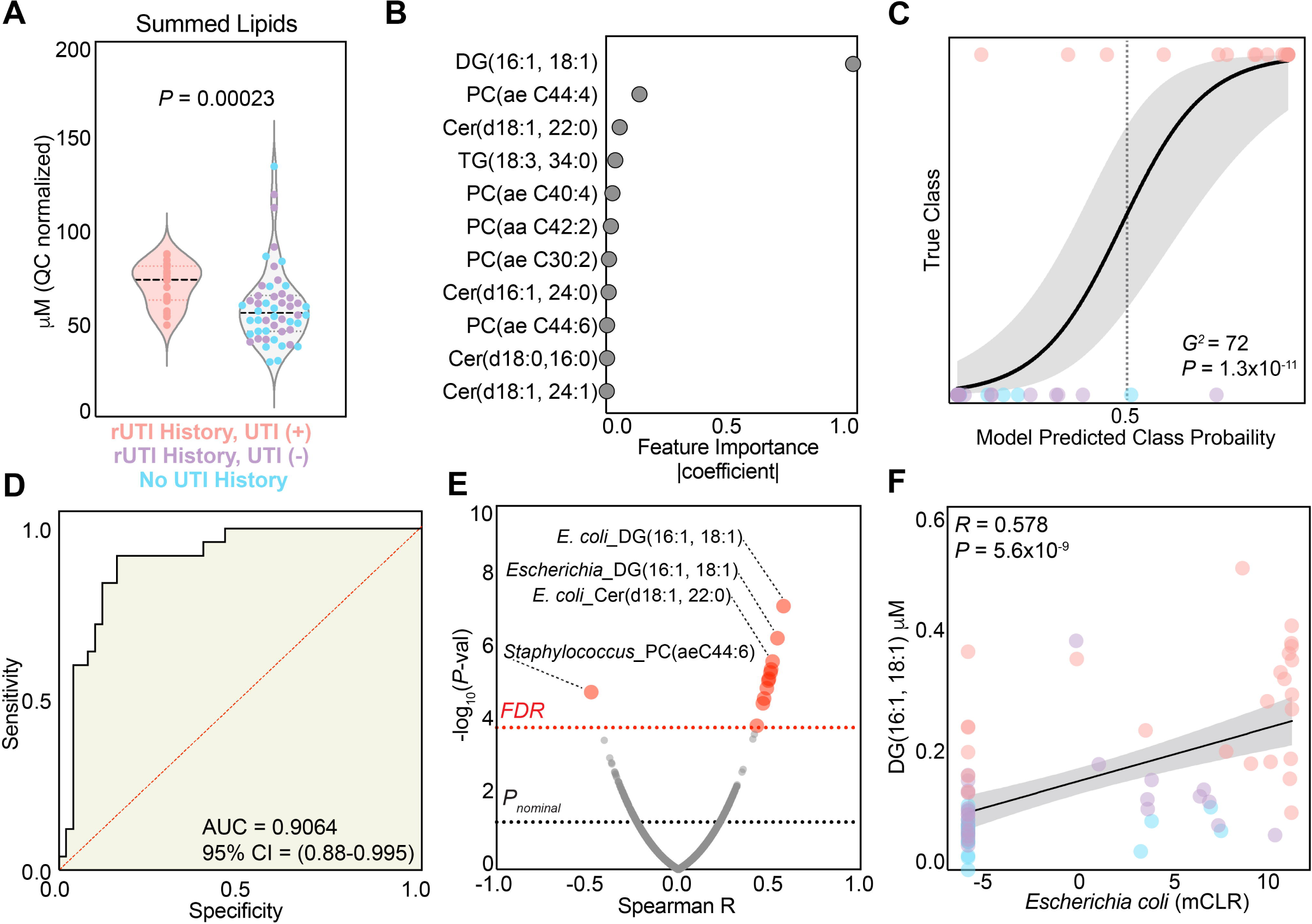
Urinary lipids as biomarkers for active rUTI. (A) Comparison of the distribution of summed lipids between the combined rUTI History, UTI(-) and No UTI History group (*n*=50) and rUTI History, UTI(+) (*n*=25) group. *P-*value generated by the Wilcoxon Rank Sum test. (B) Importance of lipid features to a discriminating model identified by elastic net regularization. (C) Logistic regression performed to assess the ability of the 11 lipids to discriminate case from control samples. *P*-value generated by log-rank test. (D) Receiver Operating Characteristic curve assessing the performance of the 11-lipid model in discriminating UTI from control samples. (E) Spearman correlation analysis between urinary taxa and the 11 lipids within the discriminating model. (F) Scatter plot of the correlation between diacylglycerol (16:1, 18:1) and mCLR transformed *Escherichia coli* abundance. *P*-value generated by permutation.

To assess the association of taxa abundant in women with active rUTI with the lipid signature of rUTI, we performed a correlation analysis with the sample-matched taxonomic genus and species level profile previously generated by whole genome metagenomics. The strongest taxa-lipid positive associations were observed between diacylglycerols (DGs) and *Escherichia* or *E. coli* (**Fig. 6E, Table S8**). *E. coli* abundance was strongly correlated with urinary DG(16:1, 18:1) concentration (*R* = 0.578, *P* = 5.6×10^-9^) (**Fig. 6F**). From these data, we hypothesize that the increase in urinary lipids observed in individuals with active rUTI is associated with acute tissue damage (i.e. epithelial cell death and exfoliation) that occurs during infection.^50^ These urinary lipids may be a novel source of diagnostic biomarkers for active UTI with rapid translatability.^51^

### The Urinary Metabolome Harbors Translatable Prognostic Indicators of rUTI

We next asked if any urinary metabolites were associated with an increased hazard of recurrent UTI. Of the 25 participants with active UTI in the rUTI History, UTI(+) (Group 3), 19 had reliable time-to-next UTI metadata. We performed Cox proportional hazard analysis to identify metabolites that were significantly associated with UTI recurrence rate. The urinary concentration of 12 metabolites was significantly associated with increased hazard of UTI recurrence within the follow-up period (**Fig. 7A; Table S9**). Interestingly, the secondary bile acid, Deoxycholic acid (DCA), was the metabolite most significantly associated with increased hazard of recurrent UTI (*HR* = 3.99, *P* = 0.0015). The remaining significant metabolites identified included three cholesterol esters (CE), Taurine, three Diacylglycerols (DG), the secondary bile acid Glycochendeoxycholic acid (GCDCA), Phosphatidylcholine (C30:2), Homoarginine, and Glycosylceramide (18:1, 18:0) which all had hazard ratios ranging between two and three (**Fig. 7B**).

**Figure 7.**
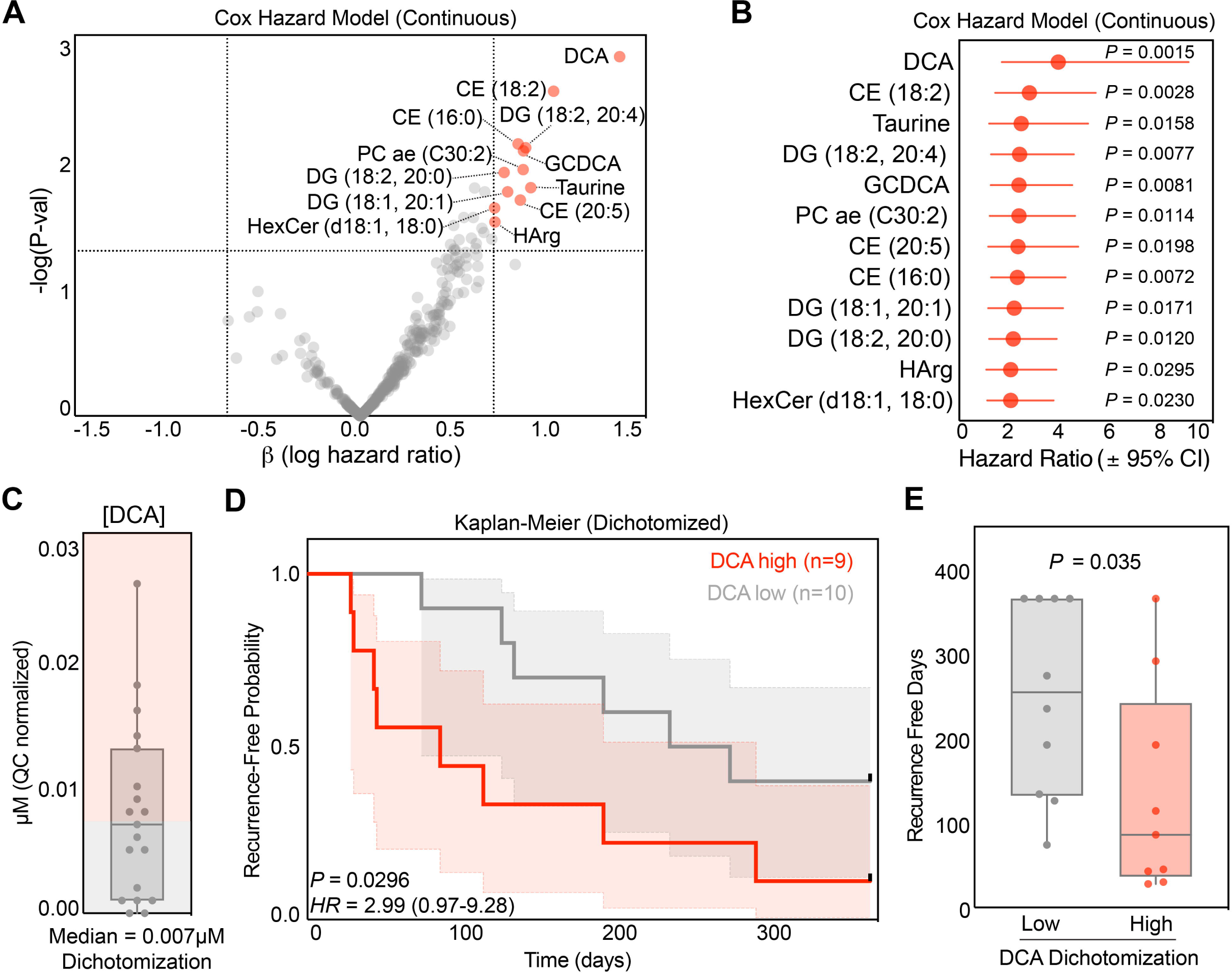
The Urinary Metabolome Harbors Translatable Prognostic Indicators of rUTI. (A) Cox proportional hazard analysis performed on metabolites on the 19 samples within the rUTI History, UTI(+) cohort that have time to recurrence metadata. (B) Hazard ratio of the 12 metabolites identified with significant association with increased hazard to recurrence. (C) DCA dichotomization scheme for Kaplan-Meier analysis. (D) Kaplan-Meier analysis of recurrence free probability and the urinary concentration of deoxycholic acid (DCA). Patients are dichotomized by the median DCA concentration. *P*-value generated by the Gehan-Breslow-Wilcoxon test. (E) Comparison of the number of Recurrence Free Survival Days between DCA low (grey) and DCA high (red) samples. *P*-value generated by Wilcoxon Rank Sum test.

To further characterize the association between urinary DCA concentration and the risk of UTI recurrence, we performed Kaplan-Meier analysis, dichotomizing the patients by the median urinary DCA concentration ([DCA] = 0.007µM, *n_high_* = 9, *n_low_* = 10) (**Fig. 7C**). Women with above median urinary concentrations of DCA exhibited a hazard ratio (HR) of 2.99 (*P* = 0.0296) for UTI recurrence compared to DCA low patients (**Fig. 7D**). Considering the differences between continuous hazard models, such as the Cox model, and dichotomous Kaplan-Meier, we consider the hazard identified by both models to be in line with one another and supportive of DCA as a prognostic indicator for rUTI. Indeed, the median recurrence-free days for women in high urinary DCA was 86, compared to 254.5 recurrence-free days in the DCA low group (*P* = 0.035) (**Fig. 7E**). While these data need to be validated in a larger patient population, they suggest that the urine metabolome may harbor valuable prognostic biomarkers for improved rUTI clinical management.

## Discussion

Here, we present a comprehensive analysis that integrates both metagenomics and metabolomics of the urinary microbial niche. We identify novel potential biomarkers of UTI and characterize how the urinary microbial and biochemical ecologies are interrelated. A critical challenge in metabolomic experiments that sample a host-microbiome ecological niche is distinguishing between host-derived and microbially derived metabolites.^52,53^ While certain metabolites can be reasonably inferred to originate from either the host or microbiome, future research will be necessary to fully untangle the origins of metabolites and broader metabolic signatures in the host-urinary microbiome relationship. Our targeted LC-MS/MS analysis provided a broad and detailed overview of the urinary metabolome, capturing both polar and non-polar metabolites. Targeted approaches offer robust estimates of absolute analyte concentrations in contrast to the relative abundance estimates typical of untargeted metabolomics approaches.^32^ However, targeted metabolomics cannot assess features not included in a predefined list of quantified targets. This limitation is significantly mitigated by the extensive range of analytes included in the Biocrates MxP® Quant 500 assay used here. Generally, we found that polar metabolites were present at higher concentrations in urine compared to the detected lipidome. This is expected due to the lower aqueous solubility of lipid species. It is important to note that, although the samples were centrifuged to remove larger particulates, no aqueous/organic partitioning was performed to separate lipids from polar metabolites. Lipid signals detected could have originated from cellular debris not pelleted by centrifugation. Given that our aim was to identify clinically translatable metabolic markers of disease and biochemical ecology, we believe that this limited sample processing most closely reflects what would be most feasible in a clinical setting.

Little is known about the urinary microbial ecosystem beyond its taxonomic profile and how it is affected by urologic disease.^9^ We aimed to characterize the urinary biochemical landscape by applying an ecological perspective to urinary metabolomic data, examining its covariance with taxonomic ecology derived from paired metagenomic data representing the same sample and microbial niche. Interestingly, no association was found between urinary biochemical α-diversity and urinary microbiome α-diversity. However, this is not completely unexpected because in this ecosystem α-diversity is not predictive of disease state alone as both UTI and a healthy urinary microbiome are often characterized by the predominance of a single or few taxa. It is also possible that dietary and lifestyle factors likely play a more significant role in shaping urinary biochemical diversity than urinary microbiome composition. However, we did identify specific metabolites linked to urinary microbiome α-diversity. Methionine sulfoxide (Met-SO), an oxidized form of methionine, was significantly correlated with urinary microbiome α-diversity. Met-SO is known to accumulate during oxidative stress and inflammation.^54,55^ Furthermore, TMAO is known to be associated with cardiovascular disease and hypertension.^56^ Given that a “healthy” urinary microbiome is often characterized by low α-diversity and predominance by *Lactobacillus* species, it is possible that Met-SO serves as a marker of inflammation associated with a more diverse urinary microbiome.

We also observed an association between the β-diversity of biochemical and taxonomic ecologies of urine, finding that as urine samples became more taxonomically distinct, they also became more metabolically distinct. However, this significant association was only observed when biochemical β-diversity was calculated using the Canberra distance. We interpret this to suggest that the metabolites driving this association are likely of low abundance as the Canberra distance normalizes individual feature distances before summing to generate the aggregate distance, which in contrast with the Bray-Curtis distance, allows analysis across a wide range of feature abundances.^57^ Based on these findings, we propose that the association between the urinary microbiome and metabolome is modest and primarily driven by the most abundant taxa and the least abundant metabolites. Given that host diet and lifestyle likely strongly influence urinary metabolite composition and that the urinary microbiome is a relatively low-biomass microbial community, we find this interpretation reasonable.^26^

Active UTI dramatically shifts the taxonomic and genomic landscape of the urinary microbiome, as we have previously shown in this same cohort.^5,23^ We hypothesized that these large-scale taxonomic shifts would also correlate with significant changes in the urinary metabolome. Our analysis uncovered not only novel metabolic enrichments associated with infection and health but also confirmed known metabolic enrichments observed in UPEC infection, such as putrescine.^28^ Interestingly, we found a substantial number of lipids enriched in samples during active UTI. Two diacylglycerols, DG(14:0, 20:0) and DG(16:1, 18:1), were the most significantly enriched metabolites in the urine of women with active UTI. Many ceramides and ceramide derivatives were also enriched in the urine of women with active UTI, which we hypothesize are a signature of bladder epithelial tissue damage as the bladder is a particularly ceramide-rich tissue and ceramide biosynthesis is not widespread among bacteria.^58,59^ The finding of a unique lipid signature associated with active UTI provides rationale for the exploration of new metabolite-based diagnostic paradigms for UTI.

The need for novel point-of-care (POC) diagnostics for UTI is significant, as current dipstick-based approaches achieve a specificity of only around 50%, and the diagnostic window for current “gold standard” culture-based methods is 24-48 hours.^36,51^ We identified a multivariate model of 11 lipids that showed outstanding performance in distinguishing active UTIs from uninfected controls. Although this finding needs to be validated in an independent cohort, it shows promise as rapid methods for lipid detection, such as matrix-assisted laser desorption/ionization time of flight (MALDI) mass spectrometry, offer opportunities to translate these biomarkers for use in clinical diagnostics.^60,61^ Furthermore, while the urinary abundance of UPEC was positively correlated with many of the lipids in this model, we also observed elevated concentrations of discriminating lipids in UTI patients without UPEC infection. Thus, this urinary lipid signature may. Have broad utility in identifying active UTI, independent of causative pathogen.

Our taxa-metabolite correlation analysis within the *No UTI History* group, focusing on metabolites enriched in healthy urine, also revealed significant associations. Notably, we found negative correlations between urinary methionine sulfoxide (Met-SO) and both the genus *Lactobacillus* and the species *L. crispatus*. This finding is in line with our additional observation the urinary Met-SO is positively associated with urinary microbiome α-diversity and suggests a decrease in oxidative stress may be associated with the predominance of beneficial *Lactobacillus*. While we cannot determine whether the urinary Met-SO is derived from the bladder or from systemic sources, its association with increased oxidative stress may indicate local inflammation.^54,55^ Interestingly, *Streptococcus anginosus*, a species linked to urogenital infections and urogenital dysbiosis, exhibited a strong positive association with urinary Met-SO.^62^ Taken together, these findings suggest a plausible link between taxonomic ecological state (healthy vs. dysbiotic) and metabolic markers of inflammation. Future work may focus on translating these findings into model systems to explore the source and implications of these associations.

There is a significant lack of molecular markers available to clinicians who aim to take a personalized approach in treating rUTI. We analyzed the association between the urinary biochemical ecology and UTI recurrence hazard to identify robust candidates for prognostic biomarkers of rUTI. Our analysis uncovered 12 candidate urinary metabolites associated with an increased hazard of UTI recurrence. The top candidate was deoxycholic acid (DCA), a microbially produced secondary bile acid, which had a hazard ratio of ∼4 in a Cox Hazard Model and ∼3 via Kaplan-Meier analysis when dichotomized by median DCA concentration.^63^ The source of DCA is particularly noteworthy. While it is possible that local microbial communities in the urinary tract are producing DCA from host-derived cholic acid, increased urinary DCA may alternatively reflect systemic gut dysbiosis, potentially affecting gut permeability, bile acid cycling, and host-microbe interactions in the gastrointestinal tract. It is conceivable that elevated circulating DCA is secreted into the urine as a result of gut dysbiosis and inflammation, which may be linked to increased susceptibility to rUTI. This is further evidenced by our observation that an additional bile acid, glycochenodeoxycholic acid (GCDCA), was also prognostic of UTI recurrence hazard. GCDCA is a host derived secondary bile acid produced in the liver and is involved in lipid uptake.^64^ This hypothesis aligns with the work of Worby et al. and Thanert et al., who contemporaneously provided evidence for gut dysbiosis, inflammation, and pathogen transmission along the gut-bladder axis in the evolution of UTI.^65,66^

Taken together our findings underscore the critical need for advancing the understanding of the urinary metabolome and its relationship with the urinary microbiome, especially in the context of rUTI and rUTI susceptibility. By identifying novel biochemical markers and elucidating metabolic associations with both microbial taxa and infection states, this work represents a meaningful step toward the development of metabolic biomarkers of UTI that can be leveraged in the development of more accurate diagnostic and prognostic paradigms. Our findings of a distinct urinary metabolome associated with microbial signatures of health and disease should be further investigated to identify metabolic interactions critical to the maintenance of a beneficial microbiota and prevention of urologic diseases like UTI. Direct interrogation of the intricate interactions between urinary biochemical and microbial ecologies will be critical to inform personalized, microbiome-based approaches to UTI treatment to ultimately improve patient outcomes and reduce UTI recurrence.

## Methods

### Cohort Curation and definitions

Human subjects were all postmenopausal females recruited from the Urology Clinic at The University of Texas Southwestern Medical Center under IRBs STU032016-006 (University of Texas Southwestern Medical Center) and 19MR001 (University of Texas at Dallas) as previously described.^5^ Table S10 summarizes clinical characteristics. Full clinical metadata can be found in Table S1 of Neugent et al. 2022.^5^ Written informed consent was collected from each patient. The following exclusion criteria were applied to screen patients for eligibility in the cohort: premenopausal or perimenopausal status; history of complicated UTI; antibiotic use within four weeks prior to urine sample collection, except in cases of culture-confirmed active infection; pelvic cancer or pelvic radiation within the last three years; current chemotherapy treatment; renal insufficiency (creatinine levels >1.5 mg/dL); post-void residual volume exceeding 100 mL; stage 3 or higher pelvic organ prolapse; pelvic surgery for incontinence within six months before urine sample collection; intermittent catheter use; neurogenic bladder; any upper urinary tract abnormalities associated with UTI; and diagnosis of Type 1 or Type 2 Diabetes Mellitus. One participant (PF21) from the rUTI History, UTI(+) group had an active, culture-confirmed UTI and had received antibiotics in the four weeks prior to sample collection. All urine samples were collected via “clean-catch” midstream methods after patient education.

### Targeted Urinary Metabolite Quantification

The Biocrates MxP® Quant 500 targeted metabolomics assay with the urine extension was used to quantitatively measure urinary metabolites.^67^ Assay was performed per manufacturer instructions on a Waters Xevo TQS tandem quadrupole mass spectrometer lined to an Acquity liquid chromatography system. The LC-MS system used for metabolomics passed the assay’s system suitability test immediately prior to data acquisition. Briefly, the Biocrates MxP® Quant 500 kit enables the quantification of up to 630 metabolites spanning 26 compound classes, including amino acids, biogenic amines, bile acids, fatty acids, phosphatidylcholines, ceramides, and di-/triglycerides. Lipids and hexoses were analyzed using flow injection analysis-tandem mass spectrometry (FIA-MS/MS), while small molecules were measured via liquid chromatography-tandem mass spectrometry (LC-MS/MS). Urine samples were slowly thawed at 4C and a 10µL aliquot was cleared of debris by centrifugation of 4000xG. Metabolite profiling was performed using a 96-well sample preparation device equipped with inserts impregnated with internal standards. The 10µL urine sample was added, followed by the addition of a phenyl isothiocyanate (PITC) solution for derivatization of specific analytes (e.g., amino acids). After derivatization, the analytes were extracted using an organic solvent and diluted before analysis. The resulting extracts were analyzed using FIA-MS/MS and LC-MS/MS with multiple reaction monitoring (MRM) for targeted detection. Metabolite concentrations were determined in Biocrates WebIDQ software for downstream analysis. We set a minimum threshold of detection in at least 7 individuals to balance stringent filtering with the need to retain biologically relevant metabolites for downstream analysis. This threshold was chosen empirically, as it provided sufficient representation across groups while minimizing the loss of metabolites that may still contribute meaningful biological insights.^68^

### Taxonomic Profiling of Metagenomic Data

Raw taxonomic profile data, generated in the original report for this cohort (PRJNA801448), was reanalyzed with the updated MetaPhLAn 4.1 database.^48^ Decontamination of potential background taxonomic signatures was performed with Decontam and analysis of the taxonomic signature identified in a water blank sample that was sequenced to observe potential environmental contaminants inevitably introduced during metagenomic DNA extraction or library preparation.^69,70^

### Analysis of Biochemical and Taxonomic ***β***-Diversity

β-diversity distances were calculated for both the Biochemical and Taxonomic profiles via the Bray-Curtis, Jaccard, Euclidean, Canberra, Kulczynski, Gower, and Manhattan methods. We then performed a screen for significant correlation between all combinations of the β-diversity distance arrays of the biochemical and taxonomic ecology. This analysis is primarily sensitive to monotonic associations that are reflected in the pairwise distance associations.^42^ We then used Procrustes of PCoA ordinations analysis over all combinations of the β-diversity distances calculations as a more sensitive analysis to identify associations between biochemical and taxonomic ecology. For all Procrustes analyses, metabolite β-diversity was used as the target while taxonomic β-diversity was rotated and scaled. PCoA was performed over 4 dimensions for both metabolite and taxonomic ecology. All ecological analysis was performed in the R statistical coding language using the vegan package.

### Network Analysis of Metabolite-Taxon Association

All-against-all Spearman correlation analysis was performed using the metabolite concentration and taxonomic relative abundance in the 50 samples from the No UTI History and rUTI History, UTI (+) groups. The modified centered log ratio (mCLR) transform was performed on the taxonomic relative abundance data to break compositionality.^71^ Significant associations between taxa and metabolites were defined as those reaching a Spearman coefficient > |0.3| and *P*-value < 0.05. Network visualization and analysis was performed in Cytoscape (v 3.10.2).^72^

### Cross Validation Analysis of Potential Diagnostic Biomarkers

Elastic Net regularization combined with logistic regression to classify lipidomic data was performed to identify lipid features that distinguish the combined No UTI History/rUTI History, UTI(-) samples from rUTI History, UTI(+) samples. To optimize model performance, repeated cross-validated Elastic Net was used to explore a range of alpha (from 0 to 1 in increments of 0.1) and corresponding lambda values, measuring performance with the area under the curve (AUC). The optimal model was selected based on the highest AUC, yielding the best alpha and lambda values, and the corresponding feature coefficients were extracted to identify the most important features for classification. Leave-one-out cross-validation (LOOCV) was performed using the optimal model to predict class labels. For each observation, the model was trained on the remaining data, and probabilities of class membership were predicted for the left-out observation. The predicted probabilities were converted to class labels based on a threshold of 0.5. Model performance was assessed using ROC analysis, with the AUC value calculated for the LOOCV predictions.

### Putative Prognostic Biomarker Analysis

To assess the association between individual metabolites and time to recurrence, a Cox proportional hazards regression analysis was performed for each metabolite in the dataset. The dataset was first loaded from a CSV file, and time to recurrence along with recurrence status were extracted as the response variables. The analysis focused on the relationship between each metabolite and the survival outcome, excluding missing values where applicable. A Cox proportional hazards model was fitted for each metabolite using the ‘surv()’ function, which models time to recurrence and recurrence status.

Kaplan-Meier survival analysis was performed to further explore the association between selected metabolites and time to recurrence. Significant metabolites of interest identified by the COX proportional hazard analysis were used, and for each metabolite, the data was dichotomized based on the median concentration. For each metabolite, values above the median were classified as “High” and values below or equal to the median were classified as “Low.” The Kaplan-Meier estimator was then applied to assess differences in survival between the high and low groups. The survival curves were generated using the ‘survfit()’ function, and the resulting Kaplan-Meier plots were visualized with 95% confidence intervals.

### Statistical Analysis

All statistical analysis was performed using the R statistical programing language (V4.4.0). GraphPad Prism 10 was used for plotting graphs and Microsoft Excel was used for data interrogation and filtering. Pairwise hypothesis testing was performed using the non-parametric Wilcoxon Rank Sum test while the Kruskal-Wallis nonparametric ANOVA with Dunn’s multiple comparison post-hoc was performed for analysis of 3 or more groups. Adjustment for multiple hypothesis testing was performed using the false discovery rate (FDR) when appropriate. We evaluated significance using both nominal and adjusted p-values. When nominal significance was applied, we provided complementary visual assessments, such as scatter plots or Kaplan-Meier plots, to support the interpretation of significance.^73^

### Data Availability

The whole genome metagenomic sequencing data (FASTQ files) were previously generated^5^ and have been deposited in the NIH Sequence Read Archive (SRA) under BioProject number PRJNA801448. Prior to deposition, all human-mapping reads were removed to ensure compliance with privacy regulations. Additionally, all R code used for data analysis is available on GitHub at https://github.com/deniscolab/Neugent_etal_Urine_Biochem_Ecology, ensuring reproducibility and transparency in our analytical methods.

## Acknowledgments

We wish to acknowledge and sincerely thank the patients who participated in this study. This work was supported by grants from the Foundation for Women’s Wellness (FWW), and the National Institutes of Health (1R01DK131267-01) to N.J.D., and (1F32DK128975-01) to M.L.N. This work was also supported by the Felicia and John Cain Distinguished Chair in Women’s Health to P.E.Z. We would like to thank all the members of the De Nisco and Shulaev labs for their creative input throughout this study. We would further like to acknowledge and thank the Genome Center at the University of Texas at Dallas for their invaluable work and assistance in generating the metagenomic dataset used for this study.

## Author Contributions

Conceptualization, M.L.N., V.S., and N.J.D.; data curation, M.L.N., N.V.H., D.G., C.N.S.; formal analysis, M.L.N; funding acquisition, M.L.N., P.E.Z., and N.J.D.; investigation, M.L.N., N.V.H., V.S.; methodology, M.L.N., N.V.H, P.E.Z., V.S, and N.J.D.; project administration, M.L.N. and N.J.D.; resources, P.E.Z., V.S. and N.J.D.; software, M.L.N.; supervision, V.S., and N.J.D.; validation, M.L.N., N.V.H.; visualization, M.L.N., and N.J.D.; writing – original draft, M.L.N., and N.J.D.

