## Supplementary Data for "Urinary biochemical ecology reveals microbiome-metabolite interactions and metabolic markers of recurrent urinary tract infection"

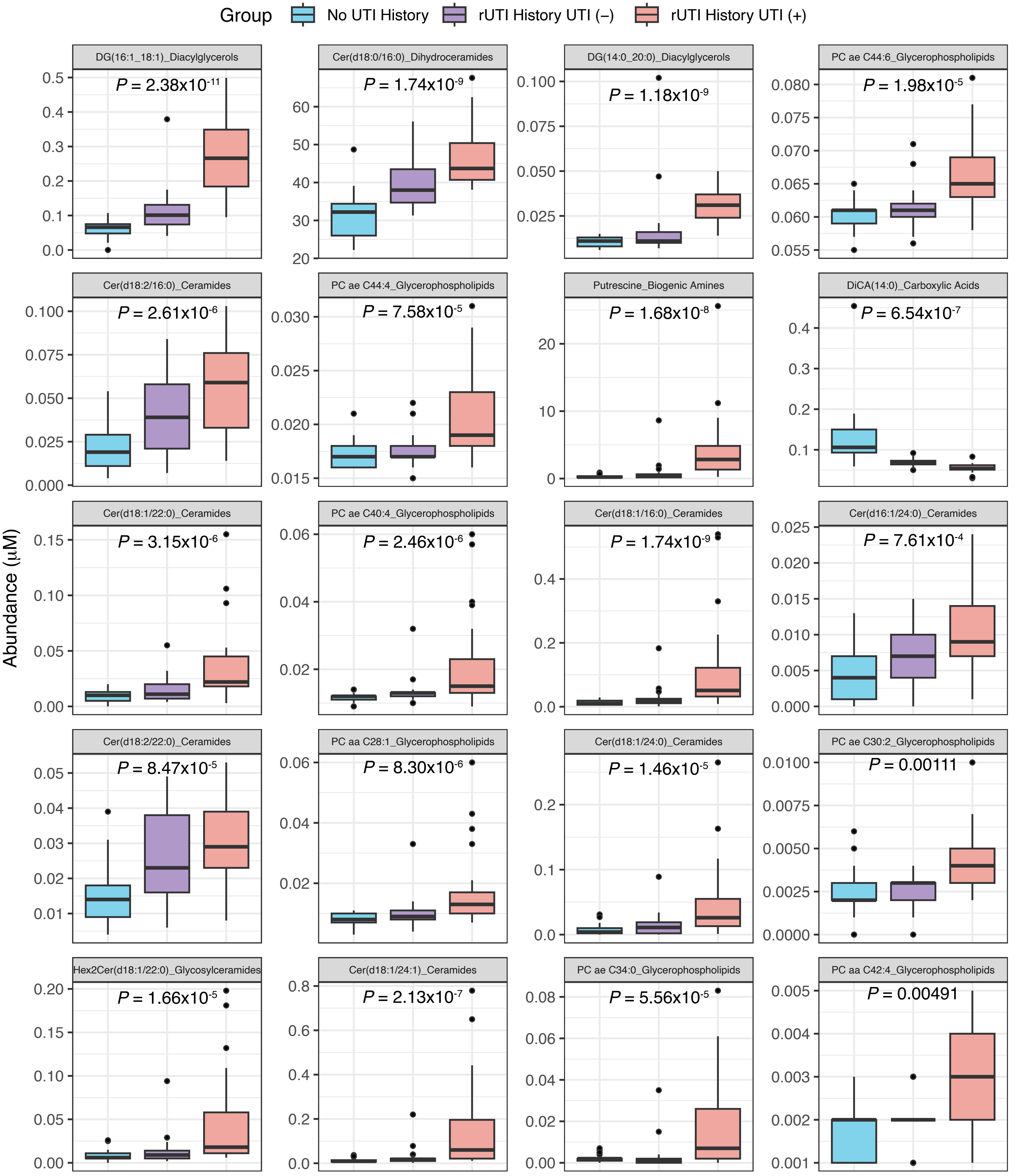


**Figure S1.** Top 20 metabolites associated with group identity. Box plots of the top 20 Variable Importance in Projection (VIP) scoring metabolites identified by PLS-DA among the three groups in the cohort studied. P-value generated by Kruskal-Wallis non-parametric ANOVA.

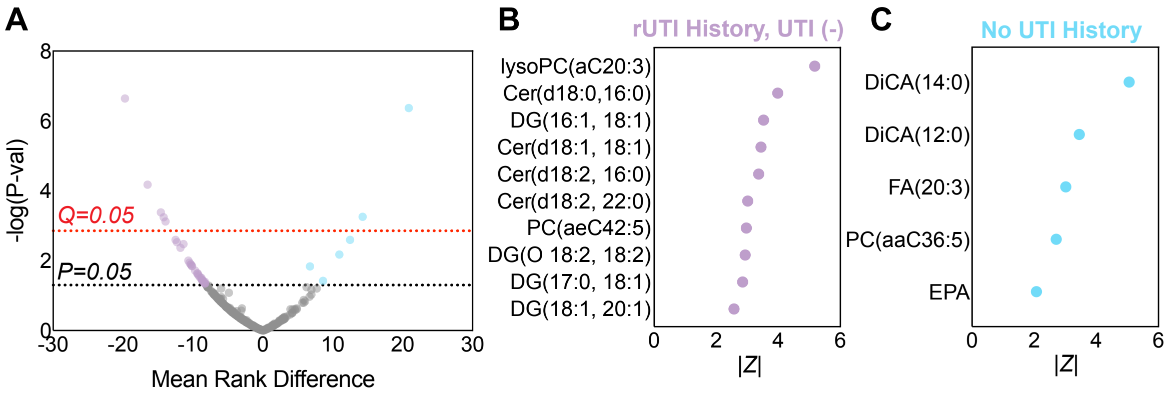


**Figure S2.** Differential Metabolite Enrichment Between No UTI History and rUTI History, UTI (-) groups. (A) Differential enrichment analysis of urinary metabolites between the No UTI History (Blue) and rUTI History, UTI(-) (Purple) groups. P-value generated by the Wilcoxon Rank Sum test. Q-value generated by false discovery rate. Absolute value of the Wilcoxon Rank statistic of the top 10 metabolites differentially enriched in the (B) rUTI History, UTI (-) group and in the (C) No UTI History group.

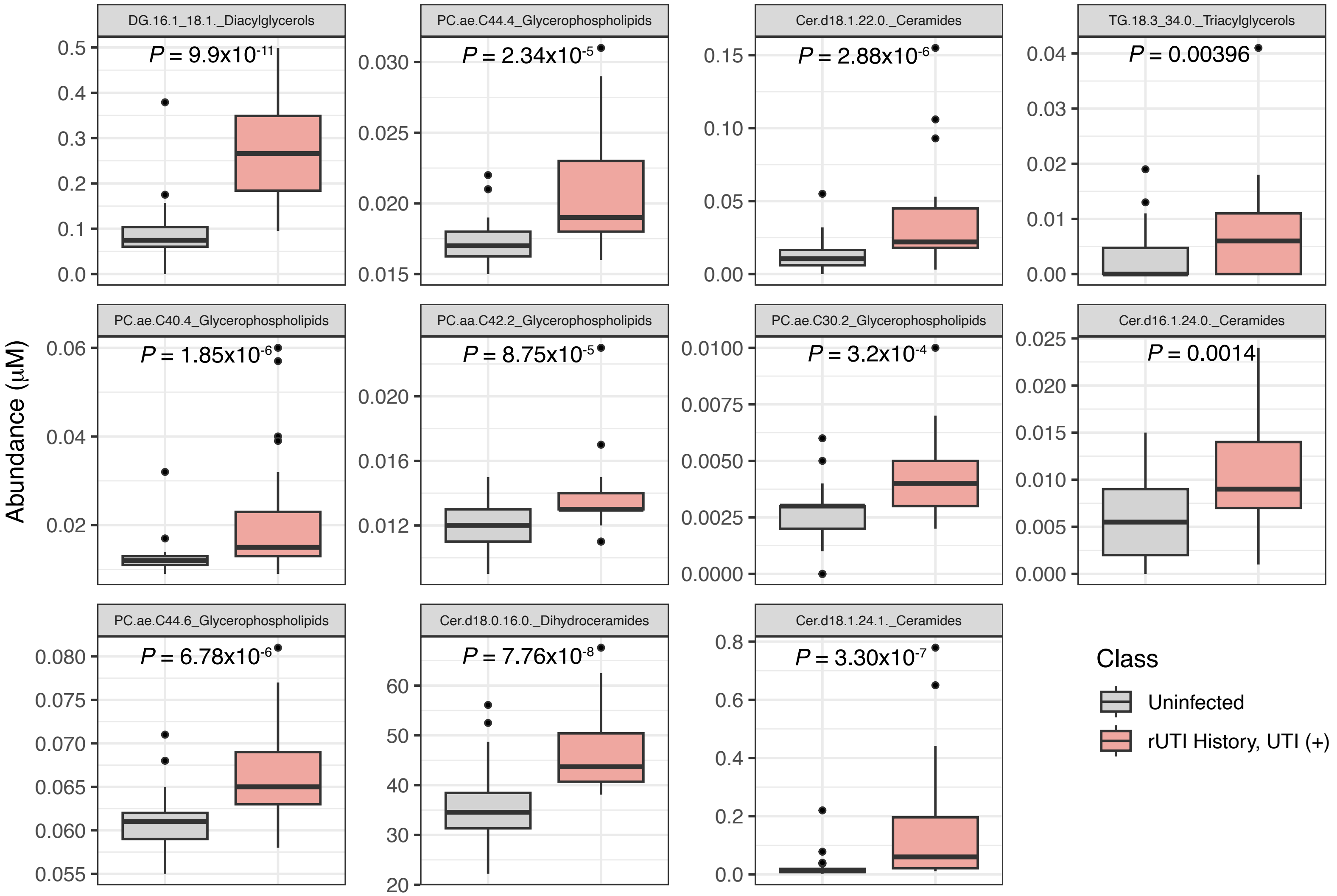


**Figure S3.** Optimal Lipids to Distinguish Uninfected from rUTI History, UTI (+) samples. Box plots of the 11 lipids identified by elastic net regularization to distinguish the combined Uninfected samples (No UTI History and rUTI History, UTI(-)) from the rUTI History, UTI (+) samples. P-value generated by Wilcoxson Rank-Sum.

| | No UTI History (Group 1) | rUTI History, UTI(-) (Group 2) | rUTI History, UTI(+) (Group 3) | *P*-value |
|---|---|---|---|---|
| ***N*** **(Individuals)** | 25 | 25 | 25 | **-** |
| **Age (years) (range)** | 67 (51-82) | 68 (52-86) | 76 (56-88) | **0.04**[A] |
| **Race** | | | | |
| *African American* | 1 (4%) | 1 (4%) | 0 (0%) | |
| *Caucasian* | 24 (96%) | 23 (92%) | 22 (88%) | |
| *Hispanic* | 0 (0%) | 1 (4%) | 3 (12%) | |
| *Other* | 0 (0%) | 0 (0%) | 0 (0%) | 0.33[B] |
| **BMI (range)** | 26.2 (19.0-34.2) | 25.3 (19.7-33.5) | 27.3 (18.7-41.6) | 0.25[A] |
| **Smoking History** | | | | |
| *Never* | 16 (64%) | 17 (68%) | 17 (68%) | |
| *Ever* | 9 (36%) | 8 (32%) | 8 (32%) | 0.94[B] |
| **EHT** | | | | |
| *EHT (-)* | 10 (40%) | 11 (44%) | 17 (68%) | |
| *EHT (+)* | 15 (60%) | 14 (56%) | 8 (32%) | 0.10[B] |
| **Urine pH (range)** | 6.0 (5.0-8.5) | 6.0 (5.0-7.5) | 5.5 (5.0-9.0) | 0.60[A] |

[A] Kruskal-Wallis test
[B] $\chi^2$ test

**Table S10: Cohort clinical characteristics**
